# Fast and ultra-sensitive glycoform analysis by supercritical fluid chromatography-tandem mass spectrometry

**DOI:** 10.1101/2021.09.30.462681

**Authors:** Yoshimi Haga, Masaki Yamada, Risa Fujii, Naomi Saichi, Takashi Yokokawa, Toshihiro Hama, Yoshihiro Hayakawa, Koji Ueda

## Abstract

Therapeutic monoclonal antibodies (mAbs) are currently the largest and fastest growing category of biopharmaceuticals. Glycosylation of mAbs has a significant impact on their effector functions, as well as on their safety and pharmacokinetics. Heterogeneity of glycoforms is affected by various factors such as the producing cells and cell culture process. Therefore, accurate glycoform characterization is essential for drug design, process optimization, manufacturing, and quality control of therapeutic mAbs. In this study, we developed a fast, quantitative, and highly sensitive analytical platform for mAb glycan profiling by supercritical fluid chromatography-tandem mass spectrometry (SFC-MS/MS). An 8-minute analysis of bevacizumab, nivolumab, ramucirumab, rituximab, and trastuzumab by SFC-MS detected a total of 102 glycoforms, with a detection limit of 5 attomole. The dynamic range of glycan abundance was over 6 orders of magnitude for bevacizumab analysis by SFC-MS, compared to 3 orders of magnitude for conventional fluorescence HPLC analysis. This method revealed the glycan profile characteristics and lot-to-lot heterogeneity of various therapeutic mAbs. We were also able to detect a series of structural variations in pharmacologically important glycan structures. SFC-MS-based glycoform profiling method will provide an ideal platform for in-depth analysis of precise glycan structure and abundance.

## Main text

Therapeutic monoclonal antibodies (mAbs) have become a powerful therapeutic modality for various diseases, including cancer, inflammatory diseases, and autoimmune diseases. By 2020, the U.S. Food and Drug Administration (FDA) has approved 84 therapeutic mAbs, and more than 500 therapeutic mAbs are being studied in clinical trials worldwide ^1^. Most of these mAbs are IgG isotypes, with two N-glycans attached to the heavy chain at Asn297 in the Fc region. Importantly, the use of mammalian expression systems results in significant heterogeneity of these glycans. Such variation of N-linked glycans can affect their biological activity, pharmacokinetics, stability, and immunogenicity of the mAbs. Therefore, it is particularly important to fully characterize the N-glycan profile and routinely monitor it to ensure the consistent quality of the therapeutic antibodies.

Due to its complex structure, the complete analysis of the N-glycan structure on glycoproteins is a very challenging task. In conventional glycan structural analysis, high-performance liquid chromatography (HPLC), capillary electrophoresis (CE), and lectin affinity chromatography are used to detect fluorescence-labeled glycans released from glycoconjugates. However, the depth of analysis, throughput, and reproducibility have been considered inadequate for the evaluation of drug quality, i.e., chemistry, manufacturing and control (CMC)^2 3^. In the last few decades, various mass spectrometry techniques have emerged as attractive alternatives to these technologies. In fact, existing mass spectrometry platforms, such as liquid chromatography (LC)-mass spectrometry (MS) and CE-MS, were able to analyze detailed glycan structures rapidly and with high sensitivity. Typically, these methods can detect glycans on the order of femtomoles (fmol) to nanomoles (nmol) within one to two hours of mass spectrometric analysis ^4 5 6^.

In order to establish a more time-efficient and sensitive glycoform analysis method for routine CMC procedures and basic research, we employed supercritical fluid chromatography (SFC)-MS for the first time and developed an ultra-fast and ultra-sensitive glycoform profiling technique (**Fig. 1a**). As the mobile phase of SFC, supercritical fluid carbon dioxide (CO_2_) was used in this study. CO_2_ has a low viscosity and a high diffusion coefficient comparable to that of a gas, allowing for fast separation of analytes at a much higher resolution and lower pressure than HPLC ^7^. Despite these great advantages, SFC has not been used for glycan analysis, which is due to the low solubility of high-polarity compounds to the supercritical fluid CO_2_ ^8^. In other words, the native form of glycans is almost insoluble in supercritical fluid CO_2_ due to its high polarity. To overcome this issue, we have established a one-pot, cleanup-free protocol for peracetylation of glycans (**Fig. 1b**). This method can replace all polar hydroxyl groups in the released N-glycans with non-polar acetyl groups. Regardless of the glycan structure, the labeling efficiency was higher than 95% (**Supplementary Figs. 1-2**). Eventually, the peracetylation pretreatment successfully enabled us to solubilize glycans in supercritical fluid CO_2_ and analyze them with SFC-MS.

**Figure 1.**
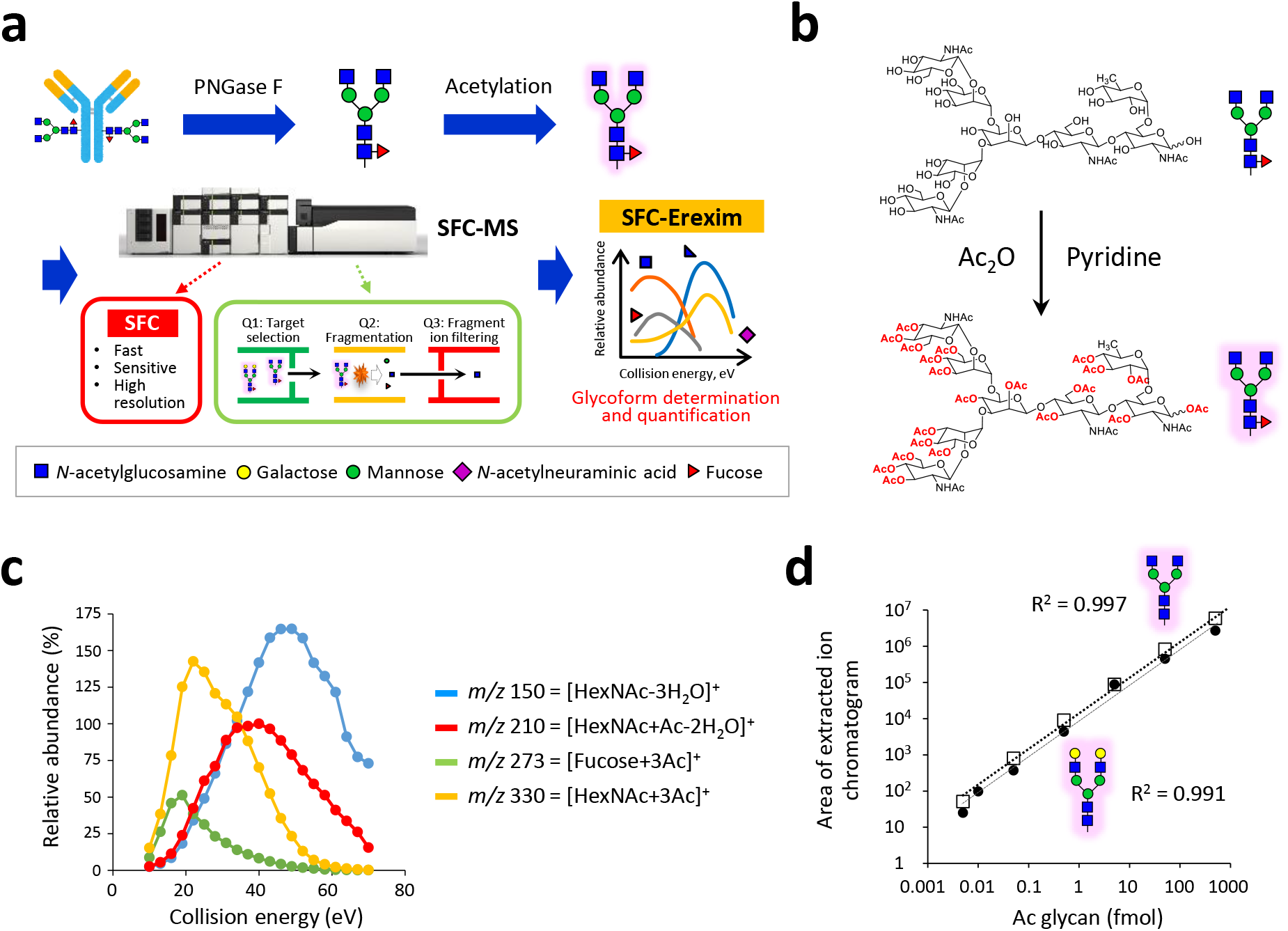
Novel glycoform profiling using SFC-MS. **a** A scheme of mass spectrometric glycan structure analysis by SFC-Erexim (energy resolved oxonium ion monitoring) technology. N-glycans are released from glycoprotein sample (immunoglobulin G), acetylated, and directly subjected to SFC-MS analysis. In the quadrupole mass spectrometer, the first quadrupole (Q1) isolated the glycan ions with unique intact mass, which are subsequently guided into the second quadrupole (Q2) where they undergo collision-induced dissociation, with continuously changed collision energies. The third quadrupole (Q3) filters the product ions to selectively detect oligosaccharide-derived ions. **b** Reaction scheme for peracetylation of N-glycans. **c** The energy-resolved product ion profiles of representative per-acetylated N-glycan (Glycan ID: 43100, the most abundant glycoform in IgG). **d** A quantitative linearity evaluation for two N-glycan standards (Glycan ID 43000 and 45000). Correlation coefficients between experimentally measured concentrations and expected ones were shown. Error bars represent the standard deviation (n = 6, independent technical experiments).

Structural characterization and quantification was performed with our energy-resolved oxonium ion monitoring (Erexim) technology, which is a kind of multiple reaction monitoring (MRM) method ^9 10^. Collision induced dissociation (CID) fragmentation of glycans/glycopeptides generates glycan-derived fragment ions, called oxonium ions ^11^. We have previously found that the original glycan structure can be determined by continuously changing the collision energy (CE) and monitoring the variety and quantity of oxonium ions. Based on this Erexim technology, we first observed the oxonium ions generated from acetylated glycans (**Supplementary Fig. 3**). Exact masses and compositions of acetylated glycan-derived oxonium ions are listed in **Supplementary Table 1**. As in the usual Erexim method, we scanned oxonium ions at various collision energy and plotted the ion yield against the collision energy (**Fig. 1c** and **Supplementary Fig. 4**). Figure 1c shows the SFC-Erexim curves obtained from the glycan composed of 4[*N*-acetylhexosamine (HexNAc)]-3[Hexose (Hex)]-1[Fucose (Fuc)]-0[*N*-acetylneuraminic acid (Neu5Ac)]-0[*N*-glycolyllneuraminic acid (Neu5Gc)] (Glycan ID: 43100). The oxonium ion *m/z* 210 derived from acetylated *N*-acetylglucosamine (GlcNAc) at the reducing end was utilized as a reporter ion for quantification. The CE for detection of *m/z* 210 was scanned and adjusted so that the intensity of this oxonium ion achieved at the maximum (**Supplementary Fig. 5**), since we have previously reported that the amount of oxonium ion produced from GlcNAc at the reducing end in this CE quantitatively correlates with the concentration of the original glycan, regardless of the glycan structure ^9^. The optimized parameters for the SFC-Erexim analysis are listed in **Supplementary Table 2**. Acetylated glycans were detected as H^+^- and/or NH_4_^+^-adduct forms in SFC-MS analysis.

The absolute sensitivity of the SFC-MS method was assessed by analyzing the peracetylated glycan standards, Glycan IDs: 43000 and 45000. The lower limit of detection (LOD) and lower limit of quantification (LOQ) were 5 attomole (amol) and 10 amol, respectively. Quantitative dynamic range was over 5 orders of magnitude (R^2^ > 0.99), highlighting the broad dynamic range of this platform (**Fig. 1d**).

SFC-MS measurements of permethylated ^12^ or peracetylated N-glycans with the reducing end labeled ^13^ were already performed three decades ago, but their sensitivity was quite low. Although the LOD was not determined in these studies, it is expected to be sub-picogram sensitivity ^13^. In this study, the resolution and sensitivity of SFC-MS as an analytical platform have been greatly improved by the increased sensitivity of the mass spectrometer and the change of an analytical method from the global method to the MRM method for targeted analysis.

Next, we applied the SFC-Erexim method to the evaluation of glycan microheterogeneity for five therapeutic antibodies (i.e., bevacizumab, nivolumab, ramucirumab, rituximab, and trastuzumab) (**Supplementary Table 3**). In order to confirm the quantitative reliability of the SFC-Erexim method, the relative abundance of each glycan quantified by this method was statistically compared with the traditional fluorescent HPLC method. As shown in Fig. 2a, the quantitative distribution patterns were almost identical between the two methods, indicating that the outcome of SFC-Erexim analysis reflects the precise glycan profile of the therapeutic antibodies (**Fig. 2a** and **Supplementary Fig. 6**). The accuracy of the quantification was maintained even though the glycans released from the antibodies equivalent to 1000 pmol by HPLC and 5 pmol by SFC-MS were measured. These results clearly show that the SFC-Erexim method is much more sensitive than the conventional fluorescence HPLC method.

**Figure 2.**
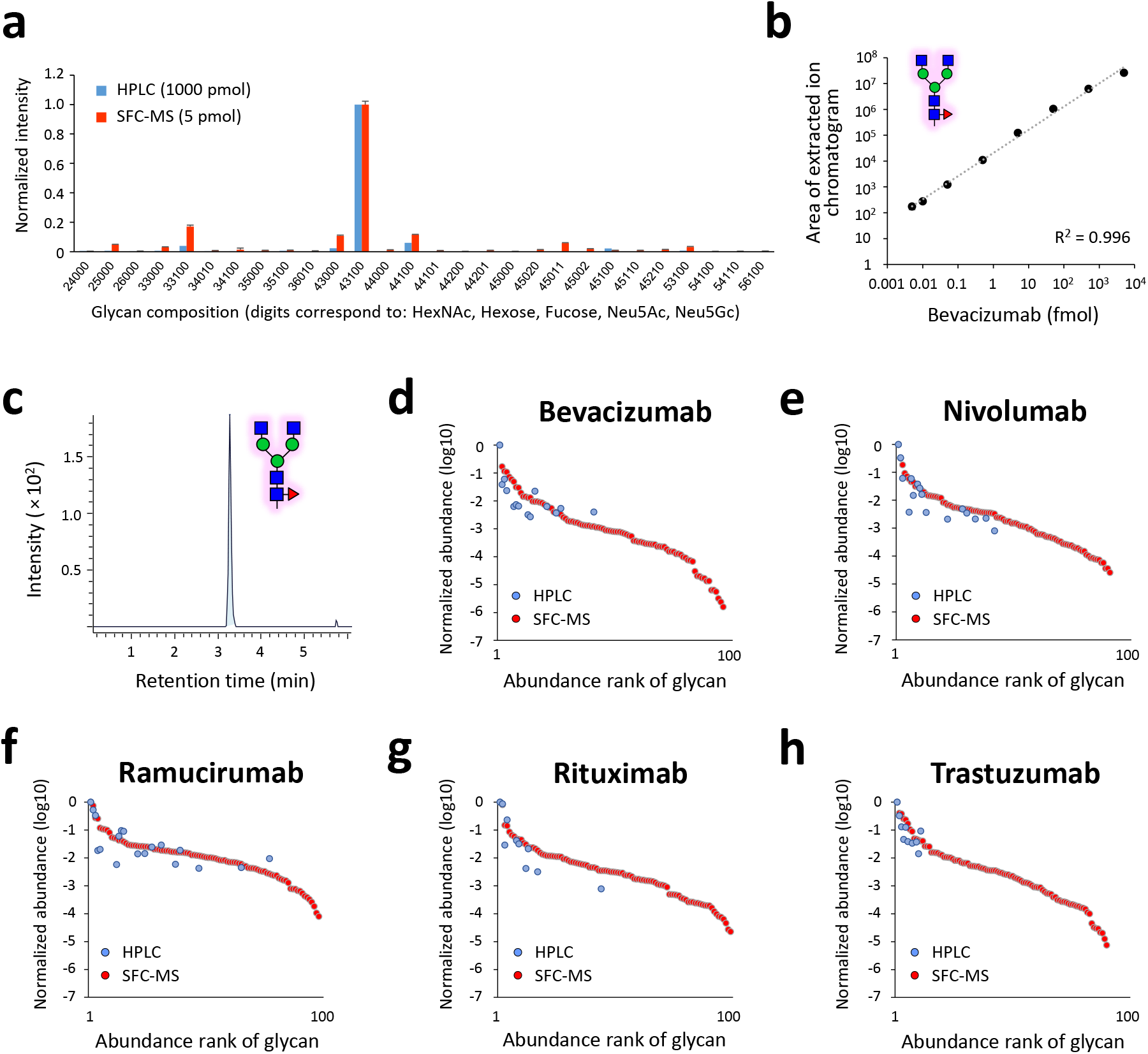
Glycoform profiling on therapeutic antibodies by SFC-MS. **a** Relative abundances of the 28 most abundant glycoforms of Bevacizumab are displayed. The released N-glycans from Bevacizumab were analyzed after PA-labeling or per-acetylation by fluorescence HPLC (1000 pmol, blue bar) or SFC-MS (5 pmol, red bar), respectively. Error bars represent the standard deviation (n =3, independent technical experiments). **b** A quantitative linearity evaluation for peracetylated N-glycan (Glycan ID: 43100, the most abundant glycoform in Bevacizumab). Correlation coefficients between experimentally measured concentrations and expected ones were shown. Error bars represent the standard deviation (n = 3, independent technical experiments). **c** Extracted ion chromatogram of 43100 glycan after injection of 10 amol of bevacizumab-derived peracetylated N-glycans. **d-h** The rank plot of detected glycoform relative abundance for (d) bevacizumab, (e) nivolumab, (f) ramucirumab, (g) rituximab and (h) trastuzumab. The released N-glycans from five therapeutic antibodies were analyzed after PA-labeling or per-acetylation by fluorescence HPLC (1000 pmol, blue) or SFC-MS (5 pmol, red), respectively.

A total of 102 glycan structures, even from the 0.1% content structures, were quantitatively detected in 8-minute SFC-Erexim runs (**Supplementary Table 4**), representing that therapeutic mAbs have huge glycan microheterogeneities. In addition, our method shows an advantage in depth of analysis compared to previous reports, which detected 21 glycoforms from bevacizumab by 60 min LC-MS run ^14^, or 19 glycoforms from rituximab and 14 glycoforms from trastuzumab by 45 min LC-MS/MS run ^15^. The lower LOD and lower LOQ of the most abundant N-glycan, Glycan ID: 43100, released from bevacizumab were 5 amol and 10 amol, respectively (**Fig. 2b-c**).

To assess the dynamic range of the SFC-Erexim method, the detected glycan structures from the five therapeutic mAbs were ranked in order of their relative abundances and displayed with measured results by the fluorescence HPLC method (**Fig. 2d-h**). These abundance plots showed that the dynamic range of the SFC-Erexim method covered 4 to 6 orders of magnitude, whereas that of the fluorescent HPLC method was 2 to 3 orders of magnitude. The number of glycan structures detected was 96 for bevacizumab by the SFC-Erexim method, compared to 15 structures by the HPLC method. Thus, the depth of our SFC-Erexim method is much greater than that of the conventional HPLC method.

The detected N-glycans were then classified according to structural characteristics and compared between the five therapeutic mAbs (**Fig. 3a** and **Supplementary Fig. 7**). Most abundant N-glycan type was neutral (i.e., not sialylated) complex-type structures for all antibodies. The relative abundance of high mannose-type glycans was found to vary widely, from 1.9% (nivolumab) to 8.4% (trastuzumab) (**Fig. 3b**). Since the high mannose-type glycans on the Fc region have been reported to significantly affect the circulating half-life of therapeutic antibodies ^16 17^, monitoring and manipulating this parameter may be valuable for controlling the pharmacokinetics of mAbs.

**Figure 3.**
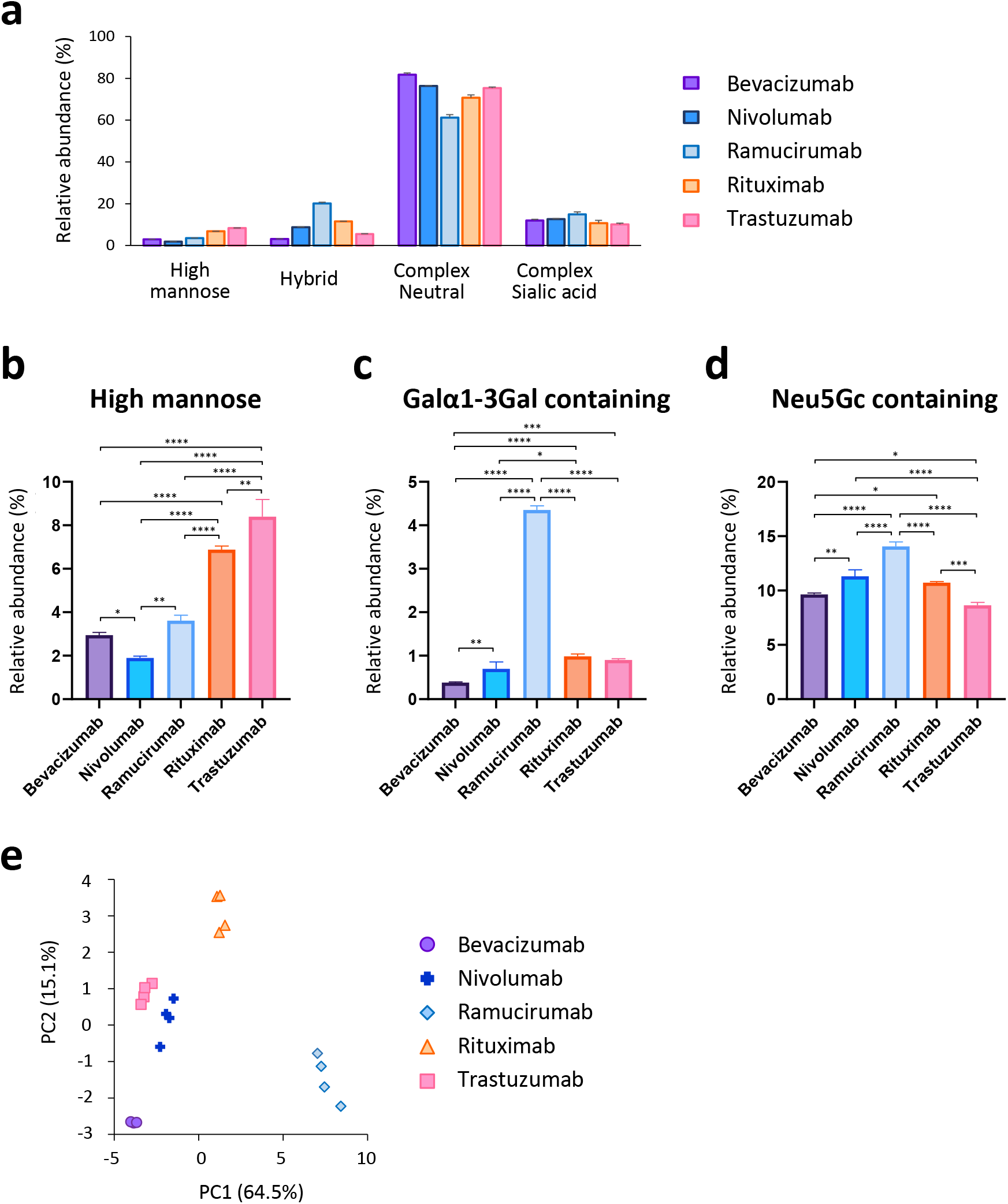
Comparison of N-glycans from five therapeutic antibodies. **a** Category comparison of N-glycome obtained from five therapeutic antibodies. Error bars represent the standard deviation determined from 3 technical replicates (for statistical analysis, see Supplementary Fig. 7). b-d Relative abundances of (b) high mannose, (c) Galα1-3Gal containing, and (d) NeuGc containing glycans of five therapeutic antibodies. Error bars represent the standard deviation (n = 3). The *p*-values were calculated by Tukey’s multiple comparisons test. Only statistically significant differences are shown. \**P* < 0.05, \*\**P* < 0.01, \*\*\**P* < 0.001, \*\*\*\**P* < 0.0001. e The principal component analysis (PCA) of five therapeutic antibodies based on N-glycome data.

Therapeutic monoclonal antibodies produced in murine myeloma cells (e.g., SP2/0 or NS0) contain terminal α1,3-linked galactose (Galα1-3Gal) structures on their Fc glycans ^18^. The Galα1-3Gal structure in the Fab region of the cetuximab heavy chain produced by SP2/0 cells has been associated with IgE-mediated anaphylactic reactions ^19 20^, however, anti-α1-3Gal IgE in allergic patients does not bind to the α1,3-galactosylated glycans on the Fc domain ^21^. Therapeutic mAbs usually lack Fab glycosylation and carry glycans only on the Fc domain ^22^, but need to be monitored and controlled for Galα1-3Gal levels during the manufacturing process. Of the five mAbs used in this study, only ramucirumab is produced in mouse NS0 cells, while the others are produced in CHO cells (**Supplementary Table 3**). Glycoform profiling clearly showed that Galα1-3Gal structures were highly expressed in ramucirumab while negligible for CHO-derived other antibodies (**Fig. 3c**). Our results also corroborated previous findings that terminal non-human sialic acid Neu5Gc are often found in in the N-glycans of recombinant IgG produced in murine myeloma and, to a much lesser extent, CHO cells ^23^ (**Fig. 3d**). It is essential to determine and control the exact proportion of these non-human immunogenic components.

To demonstrate the robustness and sensitivity of this method, we next analyzed four lots of each therapeutic antibody and evaluated lot-to-lot heterogeneity (**Supplementary Figs. 8-9**). Interestingly, among the 96 glycan structures quantified in this experiment, the abundance of the M5 isoform (high mannose-type N-glycan containing five mannose residues) showed significant lot-to-lot variation for all mAbs (**Supplementary Fig. 9**). Therapeutic IgGs containing high mannose-type N-glycans are known to be selectively cleared by a mannose receptor-mediated mechanism ^16 17 23 24^. Since high mannose-type glycans on Fc region can have a great impact on therapeutic mAb pharmacokinetics ^25^, monitoring and controlling high mannose-type N-glycan levels in manufacturing processes is important. Finally, the principal component analysis (PCA) of SFC-Erexim dataset clearly illustrated glycomic landscape-dependent classification of therapeutic mAbs (**Fig. 3e**). In this PCA plot, the glycan profile of ramucirumab was significantly different from the others, which may be due to differences in the producing cells (**Supplementary Table 3**). On the other hand, the other four mAbs were clustered separately, even though they were all produced in the same cell line, CHO. This observation indicates that the glycan profile of each therapeutic mAb is unique and is influenced not only by the type of producing cells but also by the culture conditions.

In this study, we established a highly sensitive, quantitative, and fast profiling technology for glycan structure analysis using SFC-MS. The SFC-Erexim method is considered to be a breakthrough achievement in glycan analysis because it can provide a detection sensitivity of > 5 amol within 8-min measurement with simple and low-cost procedures. Besides the antibodies targeted in this study, regulating glycosylation is very important in the development of any biologics, as glycans play an essential role in both safety and effector functions such as antibody-dependent cellular cytotoxicity (ADCC). In therapeutic mAbs alone, there are currently three commercially available glycosylated antibodies (mogamulizumab, obinutuzumab, and benralizumab) and more than 20 antibodies have been evaluated in clinical trials ^26^. Our SFC-Erexim method can also contribute to the acceleration of various innovative basic researches, such as the analysis of single-cell glycan profiles and the identification of glycan biomarkers from small amounts of clinical specimens. Based on the knowledge obtained in this report, further improvement of analytical performance and expansion of applications (e.g., O-glycosylation) are expected in the future by altering the type of supercritical fluid as a mobile phase, make-up solvent, and SFC column.

## Methods

### Reagents

Bevacizumab (Avastin^®^) and Trastuzumab (Herceptin^®^) were purchased from Chugai Pharmaceutical (Tokyo, Japan). Nivolumab (Opdivo^®^) was from Ono Pharmaceutical (Osaka, Japan), and Ramucirumab (Cyramza^®^) was from Eli Lilly Japan (Hyogo, Japan). Rituximab (Rituxan^®^) was obtained from Zenyaku Kogyo (Tokyo, Japan). Rapid PNGase F was purchased from New England Biolabs (Tokyo, Japan). BlotGlyco^®^ kit was from Sumitomo Bakelite Co. Ltd. (Tokyo, Japan), and PA-glucose oligomer was from TaKaRa (Kyoto, Japan). Sepharose CL-4B, ribonuclease B (RNase B), bovine fetuin, and LC/MS grade ammonium formate were obtained from SIGMA-Aldrich (St. Louis, MO). All other unspecified reagents were products of FUJIFILM Wako Pure Chemical Corporation (Osaka, Japan).

### N-glycan release and derivatization

N-glycans were released from proteins using Rapid PNGase F reagent according to the manufacturer’s protocol. To extract the glycans, the digest mixture was adjusted to 85% acetonitrile (ACN) (v/v). Sepharose CL-4B beads (50 μL of 50% slurry) was aliquoted onto a well of a MultiScreen Solvinert filter plate (Millipore, Billerica, MA), conditioned with water (200 μL), equilibrated with 85% (v/v) ACN (200 μL), and then loaded with sample. During the adsorption, the plate was gently shaken for 15 min at room temperature. Adsorbed samples were subsequently washed for twice in 200 μL volumes of a solution containing 0.1% trifluoroacetic acid (TFA) (v/v) in 85% (v/v) ACN, and twice with 85% (v/v) ACN. Lastly, enriched glycans were eluted in 100 μL of water. Extracted N-glycans were dried in vacuo. Released N-glycans were acetylated with 5 μl of dehydrated pyridine and 5 μl of acetic anhydride at 50°C for 4 h, and dried by SpeedVac. For HPLC analysis, released N-glycans were pyridylaminated (PA-labeling) by BlotGlyco^®^ kit according to the manufacturer’s protocol.

### SFC-MS analysis

Acetylated glycans were dissolved in methanol and analyzed using the Shimadzu Nexera UC supercritical fluid chromatography system coupled with Shimadzu LCMS-8050 triple quadrupole mass spectrometer (Shimadzu, Kyoto, Japan). CO_2_ (99.99% grade, Iwatani Corporation, Osaka, Japan) was used as the mobile phase of SFC. SFC-MS analytical conditions were as follows: modifier, methanol; injection volume, 2.5 μL; flow rate, 1.0 mL/min; make-up solvent, methanol containing 0.1% (v/v) ammonium formate; make-up flow rate, 0.1 mL/min; column oven temperature, 40°C; and column, Shim-pack UC-Phenyl (2.1 × 150 mm, 3 μm; Shimadzu, Kyoto, Japan). Gradient conditions were as follows: 10% B (0min) - 40% B (4 min) - 40% B (7 min) - 10% B (7.1 min) - stop (8.5 min). Parameters of LCMS-8050 were set as follows: interface voltage = 4000 V, interface temperature = 225°C, heat block temperature = 400°C, desolvation temperature = 225°C, CID gas pressure = 270 kPa (argon gas), Q1 resolution = unit, and Q3 resolution = unit. For relative quantification of IgG-Fc glycan microheterogeneity, the Q1 transitions were set so that they covered all possible glycan compositions. The Q3 transition was set at *m/z* 210 indicating dehydrated fragment ion of acetylated GlcNAc at the reducing terminus (i.e. [HexNAc+Ac-2H_2_O]^+^), which was utilized as a quantitative reporter ion. The CE for detection of m/z = 210 was scanned and adjusted so that intensity of this acetylated glycan-derived fragment ion achieved at the maximum (**Supplementary Fig. 5**). The optimized MRM parameters are listed in **Supplementary Table 2**.

### Fluorescent HPLC analysis

Reversed-phase HPLC was performed using an Inertsil ODS-3 column (2.1 × 150 mm, 3 μm; GL Sciences, Tokyo, Japan). The elution conditions were as follows: solvent A, 0.1 M ammonium acetate buffer, pH 4.0; solvent B, 0.1 M ammonium acetate buffer, pH 4.0, containing 0.5% 1-butanol. The column was equilibrated with 5% solvent B at a flow rate of 200 μL/min. Injected samples were separated with a linear gradient from 5 to 55% solvent B over 100 min. Fluorescence of the labeled glycans was detected at the excitation wavelength (320 nm) and the emission wavelength (400 nm). The retention times were normalized and expressed as glucose units using PA-glucose oligomer, and glycan structures were verified according to the HPLC mapping data as described previously ^27^.

### SFC-MS data processing

The acquired raw data was processed with LabSolutions LCMS software (Shimadzu) and area of product ion chromatogram for *m/z* = 210 was utilized for glycoform quantification. For relative quantification of Fc-glycan profile, the percentage composition of each glycoform was calculated with respect to the sum of all glycoforms combined. In order to exclude cross-talk between transitions where the precursor ion masses are very close, a peak area curve for correction was created and the correction value was subtracted for the corresponding glycan (**Supplementary Fig. 10**). Glycan composition is shown as Glycan ID, 5 digits correspond to: HexNAc, Hexose, Fucose, Neu5Ac, Neu5Gc.

### MALDI-TOF-MS analysis

After peracetylation, the free N-glycan were dissolved in methanol and 0.5 μL of methanol solution was applied onto a Opti-TOF™ 384-well plate (AB Sciex, Foster City, CA) together with 0.5 μL of CHCA matrix solution (4 mg/mL concentration in 70% acetonitrile, 0.1% TFA and 0.08 mg/mL ammonium citrate). The samples were left to dry by air. MALDI-TOF-MS analysis was performed using 4800 Plus MALDI TOF/TOF Analyzer (AB Sciex) operated on 4000 Series Explorer software version 3.5. The instrument was calibrated with the peptide calibration standard (AB Sciex) prior to analysis of the samples. MS measurements were performed in reflectron positive ion mode. For each spot, data was accumulated from 1000 laser shots in a randomized raster of 400 μm diameter over mass range m/z 1200-5000. The laser repeat rate was 200 Hz and the laser power was fixed at 3500 units throughout the experiment.

### Statistical analysis

Principal component analysis (PCA) was performed using Genedata Analyst software (Basel, Switzerland). Other statistical calculations for clinical data were performed using GraphPad Prism 8 software (Version 8.4.3). Intergroup differences were statistically compared by Student’s *t* test. When the means of more than 2 groups were compared, Tukey’s multiple comparisons test was used. In all these analyses, *P* < 0.05 was considered statistically significant.

## Acknowledgements

This work was supported by JSPS KAKENHI (JP19K06553) and the Project for Utilizing Glycans in the Development of Innovative Drug Discovery Technologies from the Japan Agency for Medical Research and Development (AMED).

## Author contributions

Y. Haga, M.Y., Y. Hayakawa and K.U. designed the research. Y. Haga, M.Y. and R.F. performed experiments and analyzed the data. Y. Haga and N.S. performed statistical analysis. T.Y. and T.H. provided critical reagents. Y. Haga and K.U. wrote the manuscript.

