## Supplementary Figures for "Fast and ultra-sensitive glycoform analysis by supercritical fluid chromatography-tandem mass spectrometry"

### Slide 1
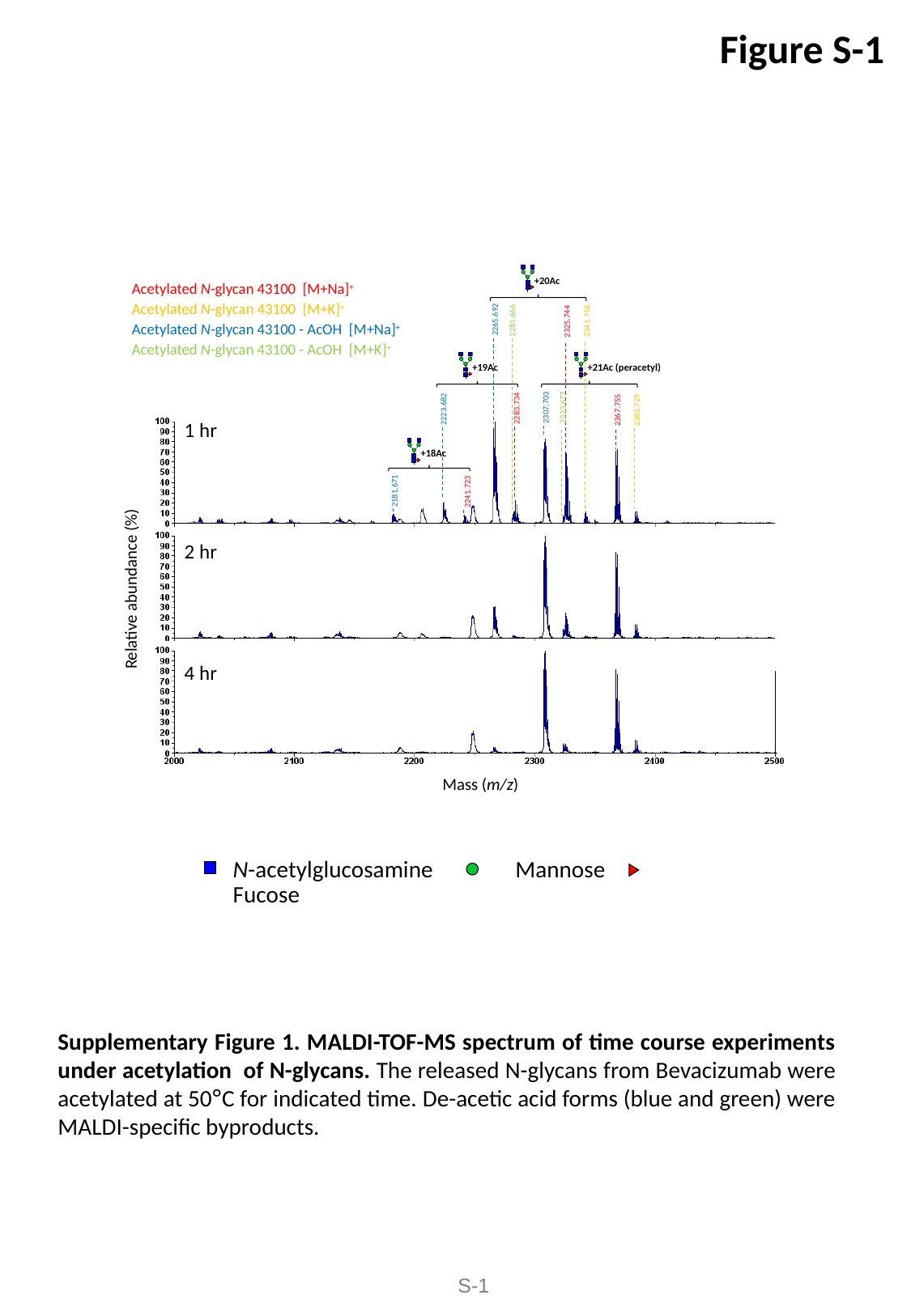

Figure S-1
+20Ac
Acetylated N-glycan 43100 [M+Na]^+^
Acetylated N-glycan 43100 [M+K]^+^
Acetylated N-glycan 43100 - AcOH [M+Na]^+^
Acetylated N-glycan 43100 - AcOH [M+K]^+^
2265.692
2281.666
2341.718
2325.744
+19Ac
+21Ac (peracetyl)
2323.677
2307.703
2283.734
2223.682
2367.755
2383.729
1 hr
2 hr
4 hr
+18Ac
2181.671
2241.723
Relative abundance (%)
Mass (m/z)
N-acetylglucosamine 　Mannose 　Fucose
Supplementary Figure 1. MALDI-TOF-MS spectrum of time course experiments under acetylation of N-glycans. The released N-glycans from Bevacizumab were acetylated at 50°C for indicated time. De-acetic acid forms (blue and green) were MALDI-specific byproducts.
S-1

### Slide 2
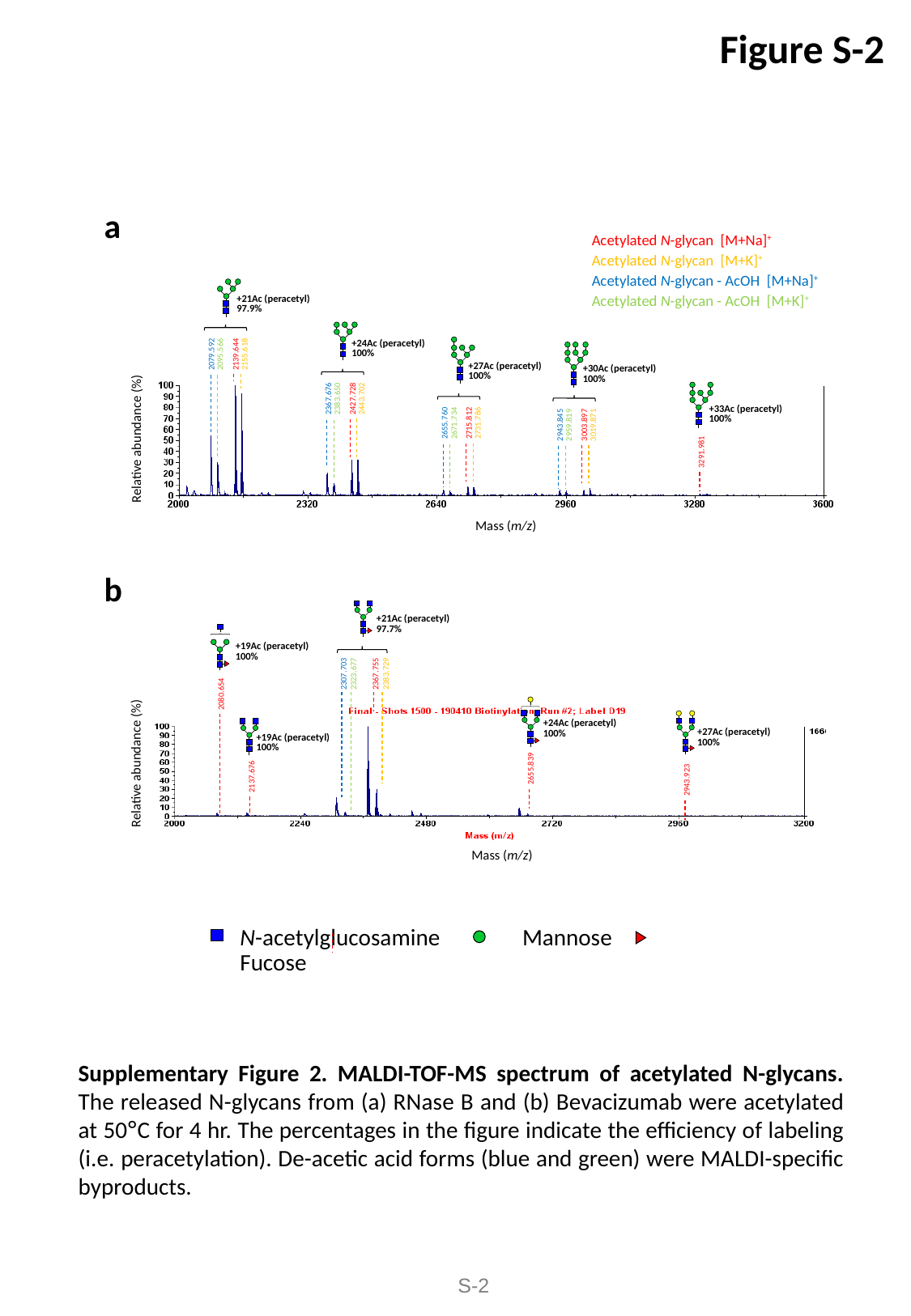

Figure S-2
a
Acetylated N-glycan [M+Na]^+^
Acetylated N-glycan [M+K]^+^
Acetylated N-glycan - AcOH [M+Na]^+^
Acetylated N-glycan - AcOH [M+K]^+^
+21Ac (peracetyl)
97.9%
+24Ac (peracetyl)
100%
2079.592
2095.566
2139.644
2155.618
Relative abundance (%)
Mass (m/z)
+27Ac (peracetyl)
100%
+30Ac (peracetyl)
100%
2367.676
2383.650
2427.728
2443.702
+33Ac (peracetyl)
100%
2655.760
2715.812
2731.786
2671.734
2943.845
2959.819
3003.897
3019.871
3291.981
b
+21Ac (peracetyl)
97.7%
+19Ac (peracetyl)
100%
2307.703
2323.677
2367.755
2383.729
Relative abundance (%)
Mass (m/z)
2080.654
+24Ac (peracetyl)
100%
+27Ac (peracetyl)
100%
+19Ac (peracetyl)
100%
2655.839
2137.676
2943.923
N-acetylglucosamine 　Mannose 　Fucose
Supplementary Figure 2. MALDI-TOF-MS spectrum of acetylated N-glycans. The released N-glycans from (a) RNase B and (b) Bevacizumab were acetylated at 50°C for 4 hr. The percentages in the figure indicate the efficiency of labeling (i.e. peracetylation). De-acetic acid forms (blue and green) were MALDI-specific byproducts.
S-2

### Slide 3
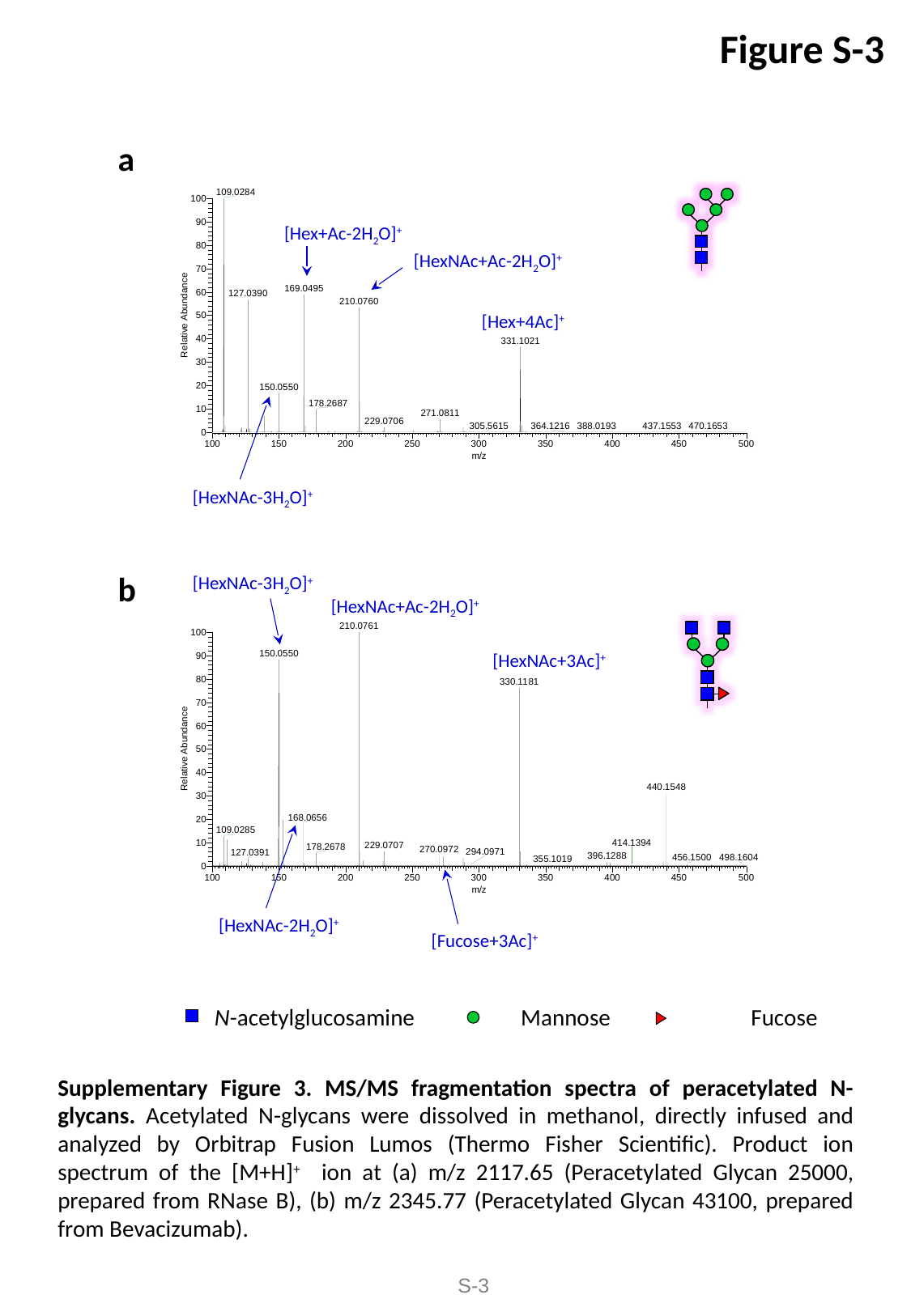

Figure S-3
a
[Hex+Ac-2H_2_O]^+^
[HexNAc+Ac-2H_2_O]^+^
[Hex+4Ac]^+^
[HexNAc-3H_2_O]^+^
[HexNAc-3H_2_O]^+^
b
[HexNAc+Ac-2H_2_O]^+^
[HexNAc+3Ac]^+^
[HexNAc-2H_2_O]^+^
[Fucose+3Ac]^+^
N-acetylglucosamine 　　Mannose 　　　Fucose
Supplementary Figure 3. MS/MS fragmentation spectra of peracetylated N-glycans. Acetylated N-glycans were dissolved in methanol, directly infused and analyzed by Orbitrap Fusion Lumos (Thermo Fisher Scientific). Product ion spectrum of the [M+H]^+^ ion at (a) m/z 2117.65 (Peracetylated Glycan 25000, prepared from RNase B), (b) m/z 2345.77 (Peracetylated Glycan 43100, prepared from Bevacizumab).
S-3

### Slide 4
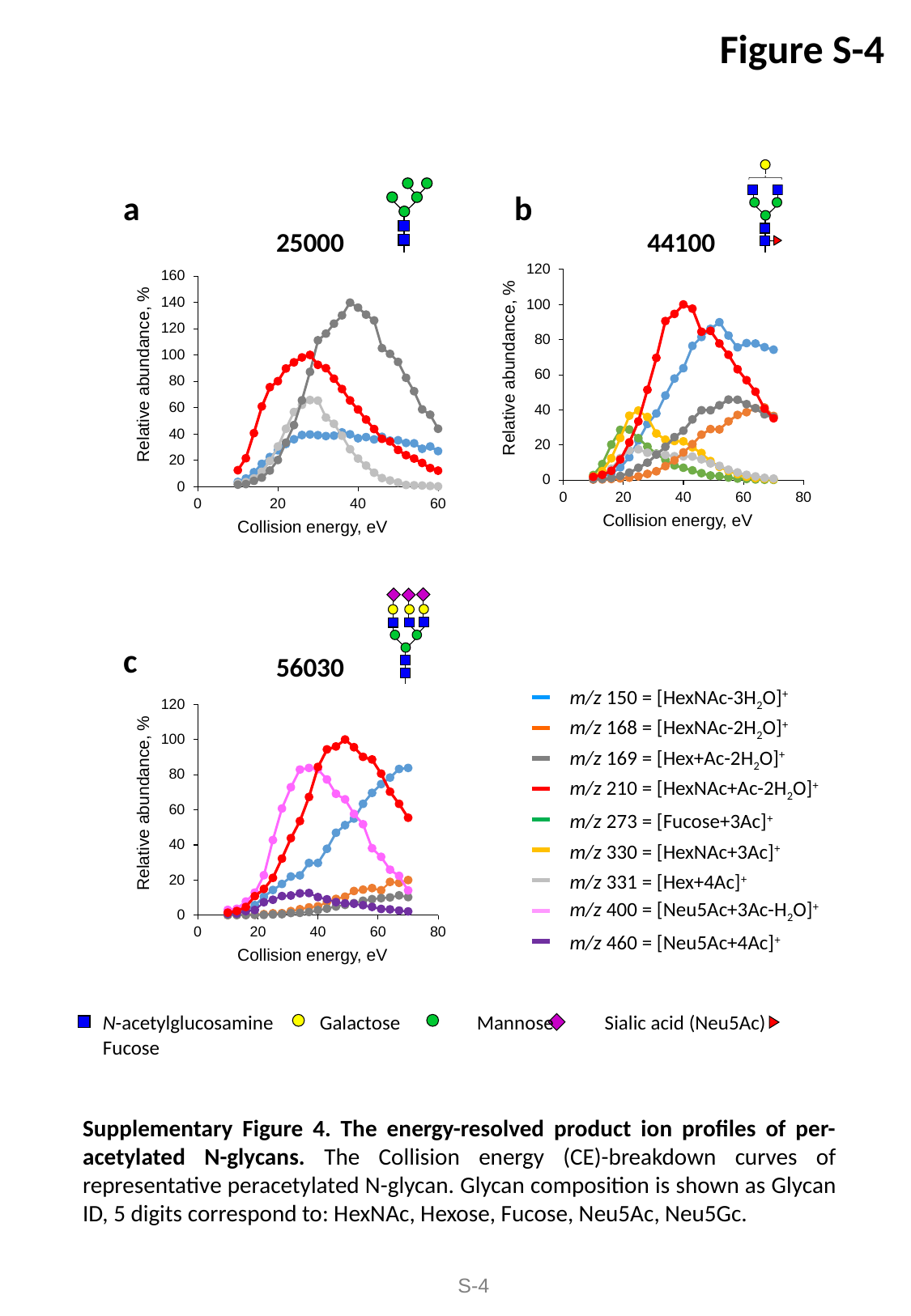

Figure S-4
b
a
25000 44100
56030
c
m/z 150 = [HexNAc-3H_2_O]^+^
m/z 168 = [HexNAc-2H_2_O]^+^
m/z 169 = [Hex+Ac-2H_2_O]^+^
m/z 210 = [HexNAc+Ac-2H_2_O]^+^
m/z 273 = [Fucose+3Ac]^+^
m/z 330 = [HexNAc+3Ac]^+^
m/z 331 = [Hex+4Ac]^+^
m/z 400 = [Neu5Ac+3Ac-H_2_O]^+^
m/z 460 = [Neu5Ac+4Ac]^+^
N-acetylglucosamine Galactose　　　 Mannose Sialic acid (Neu5Ac)　　　　Fucose
Supplementary Figure 4. The energy-resolved product ion profiles of per-acetylated N-glycans. The Collision energy (CE)-breakdown curves of representative peracetylated N-glycan. Glycan composition is shown as Glycan ID, 5 digits correspond to: HexNAc, Hexose, Fucose, Neu5Ac, Neu5Gc.
S-4

### Slide 5
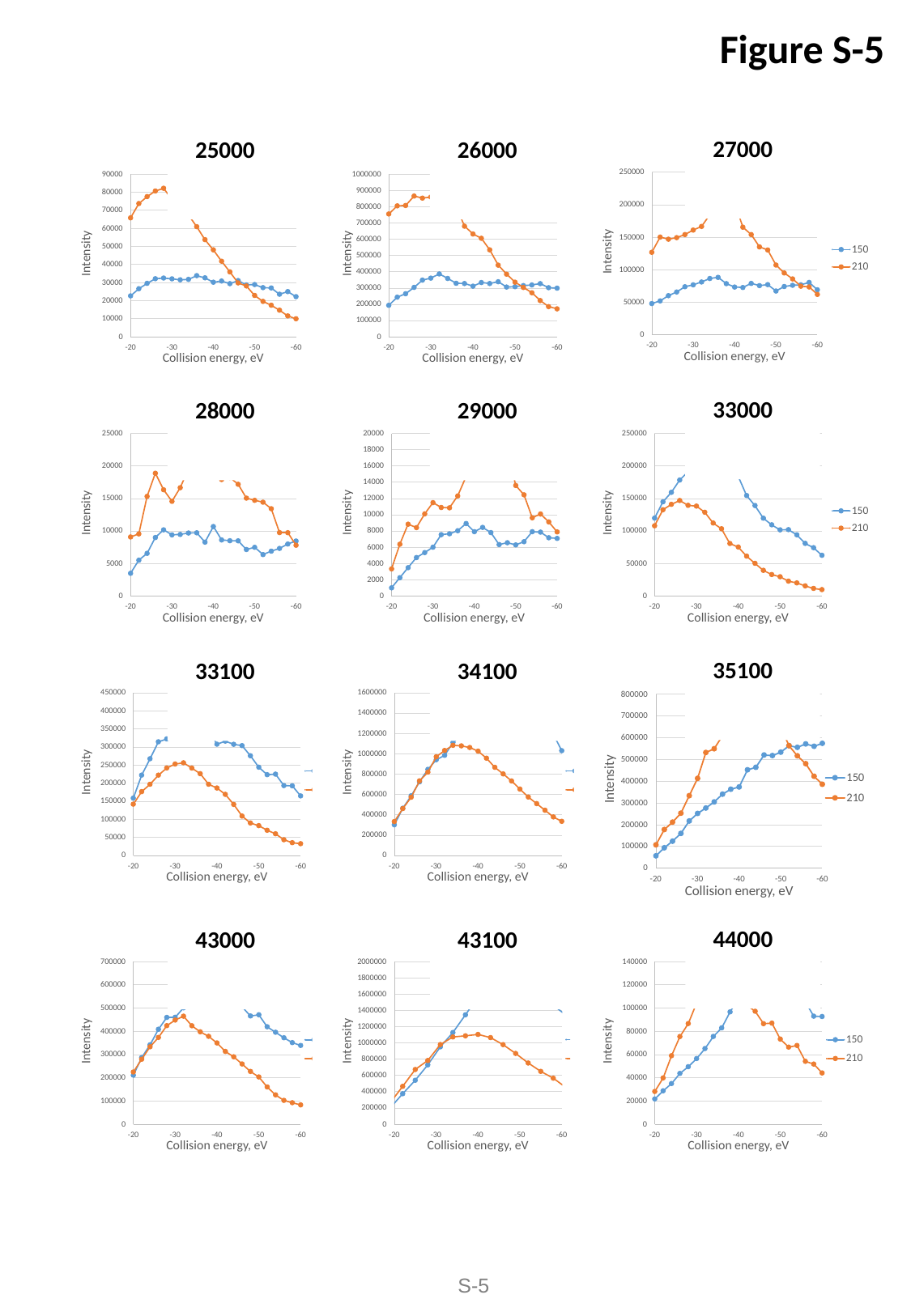

Figure S-5
 27000
 25000
 26000
 33000
 28000
 29000
 35100
 33100
 34100
 44000
 43000
 43100
S-5

### Slide 6
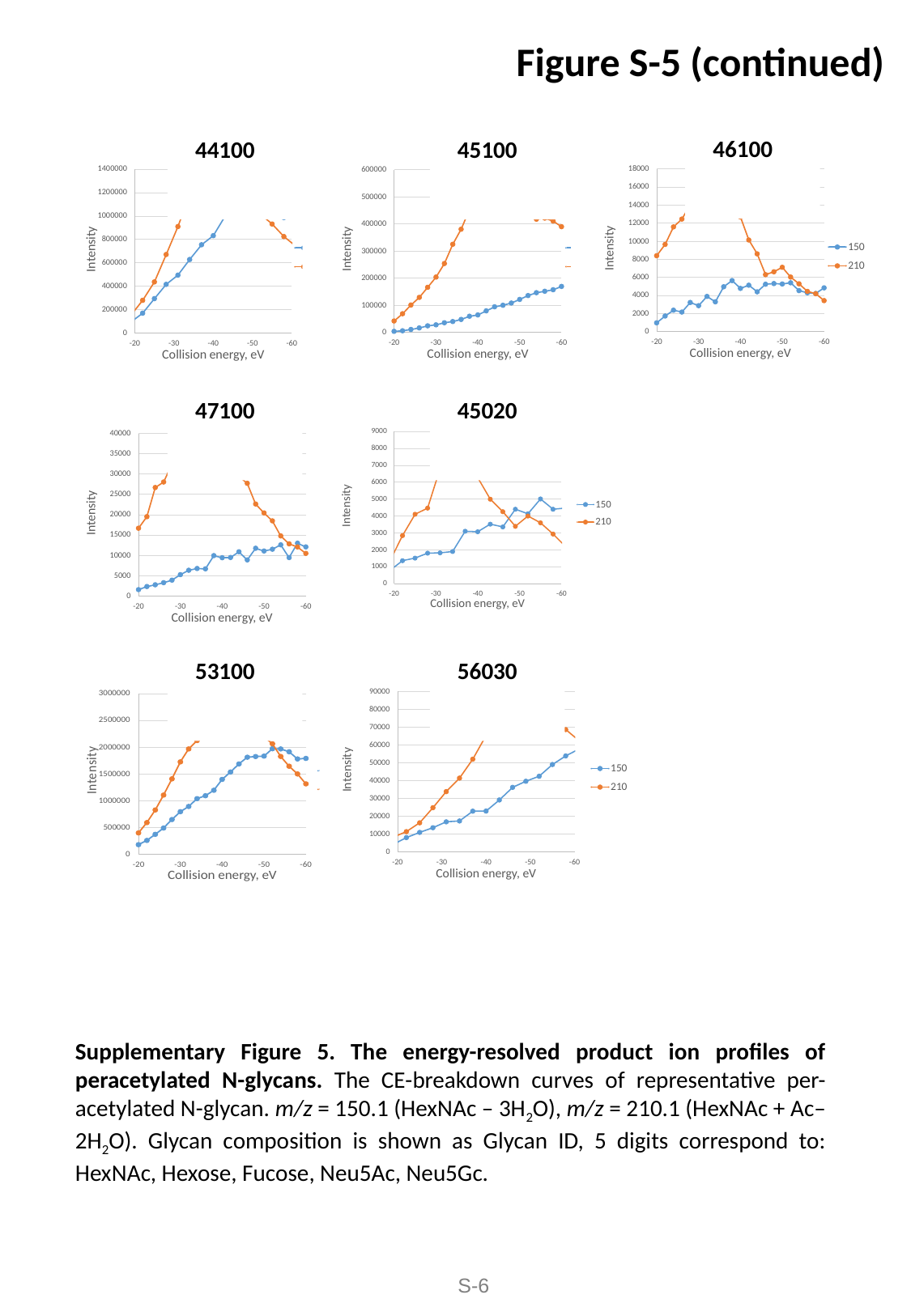

Figure S-5 (continued)
 46100
 44100
 45100
 47100
 45020
 53100
 56030
Supplementary Figure 5. The energy-resolved product ion profiles of peracetylated N-glycans. The CE-breakdown curves of representative per-acetylated N-glycan. m/z = 150.1 (HexNAc – 3H_2_O), m/z = 210.1 (HexNAc + Ac– 2H_2_O). Glycan composition is shown as Glycan ID, 5 digits correspond to: HexNAc, Hexose, Fucose, Neu5Ac, Neu5Gc.
S-6

### Slide 7
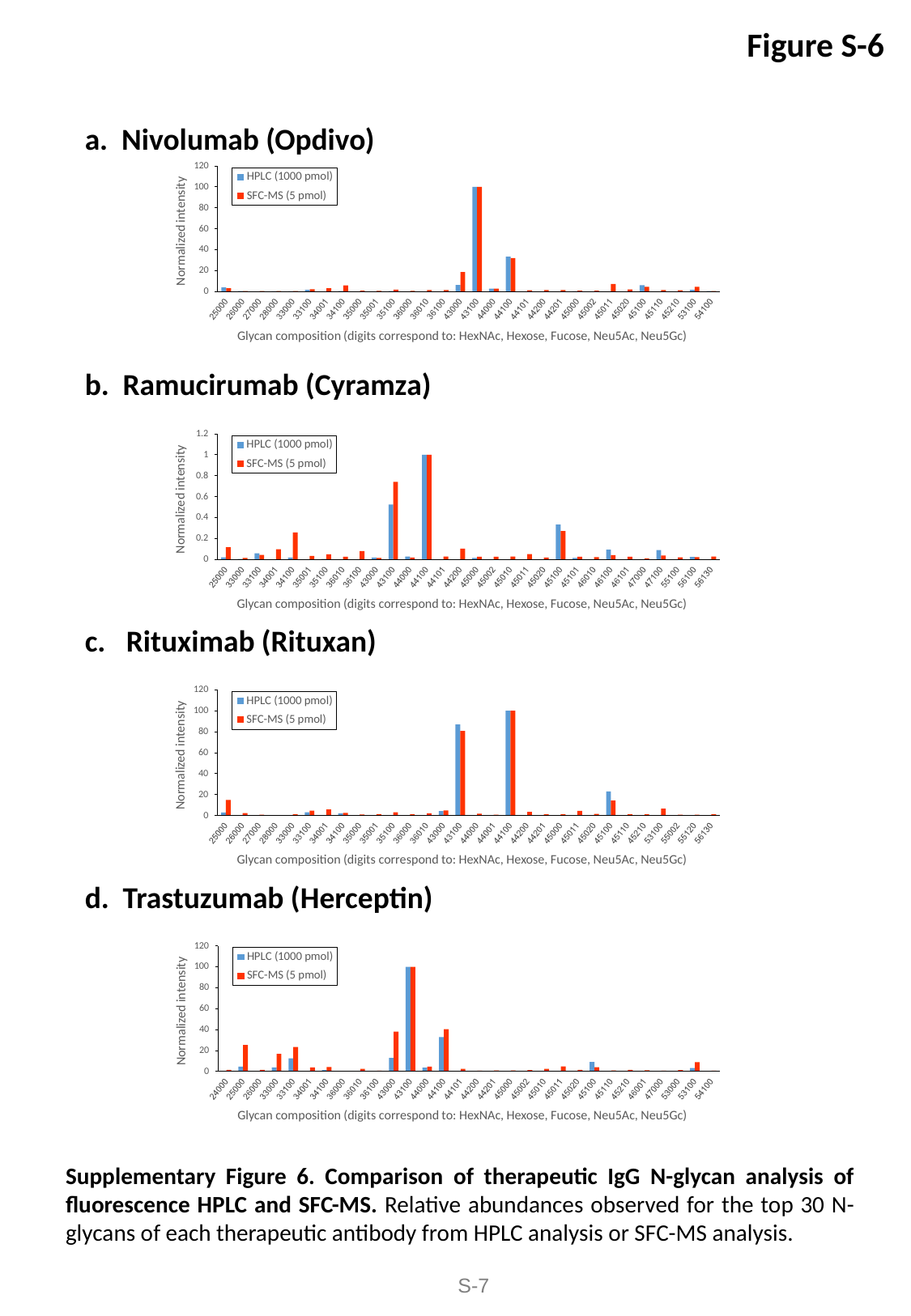

Figure S-6
a. Nivolumab (Opdivo)
b. Ramucirumab (Cyramza)
c. Rituximab (Rituxan)
d. Trastuzumab (Herceptin)
Supplementary Figure 6. Comparison of therapeutic IgG N-glycan analysis of fluorescence HPLC and SFC-MS. Relative abundances observed for the top 30 N-glycans of each therapeutic antibody from HPLC analysis or SFC-MS analysis.
S-7

### Slide 8
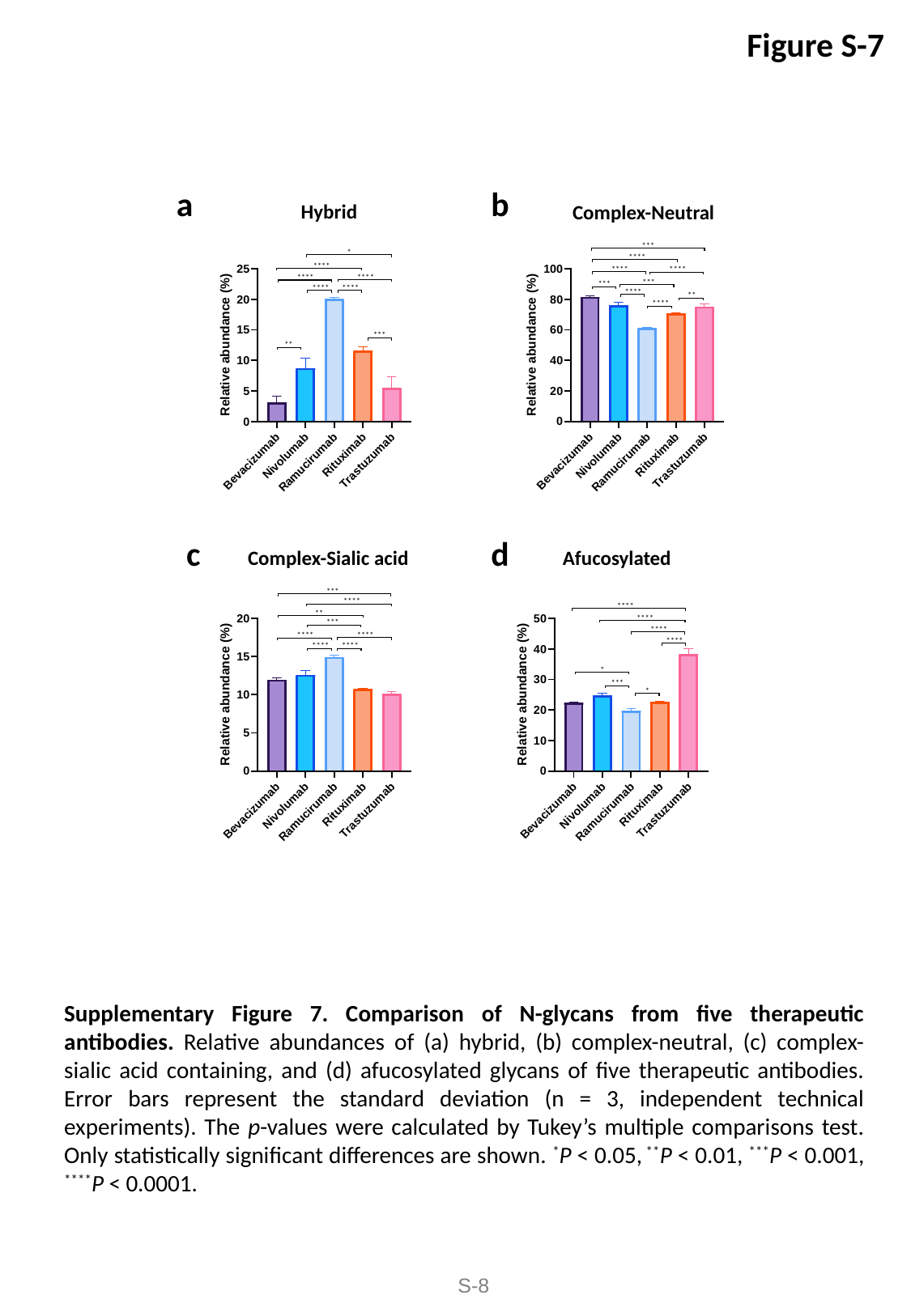

Figure S-7
a
b
Hybrid
Complex-Neutral
***
*
****
****
****
****
****
****
***
***
****
****
****
**
****
***
**
c
d
Afucosylated
Complex-Sialic acid
***
****
****
**
****
***
****
****
****
****
****
****
*
***
*
Supplementary Figure 7. Comparison of N-glycans from five therapeutic antibodies. Relative abundances of (a) hybrid, (b) complex-neutral, (c) complex-sialic acid containing, and (d) afucosylated glycans of five therapeutic antibodies. Error bars represent the standard deviation (n = 3, independent technical experiments). The p-values were calculated by Tukey’s multiple comparisons test. Only statistically significant differences are shown. ^*^P < 0.05, ^**^P < 0.01, ^***^P < 0.001, ^****^P < 0.0001.
S-8

### Slide 9
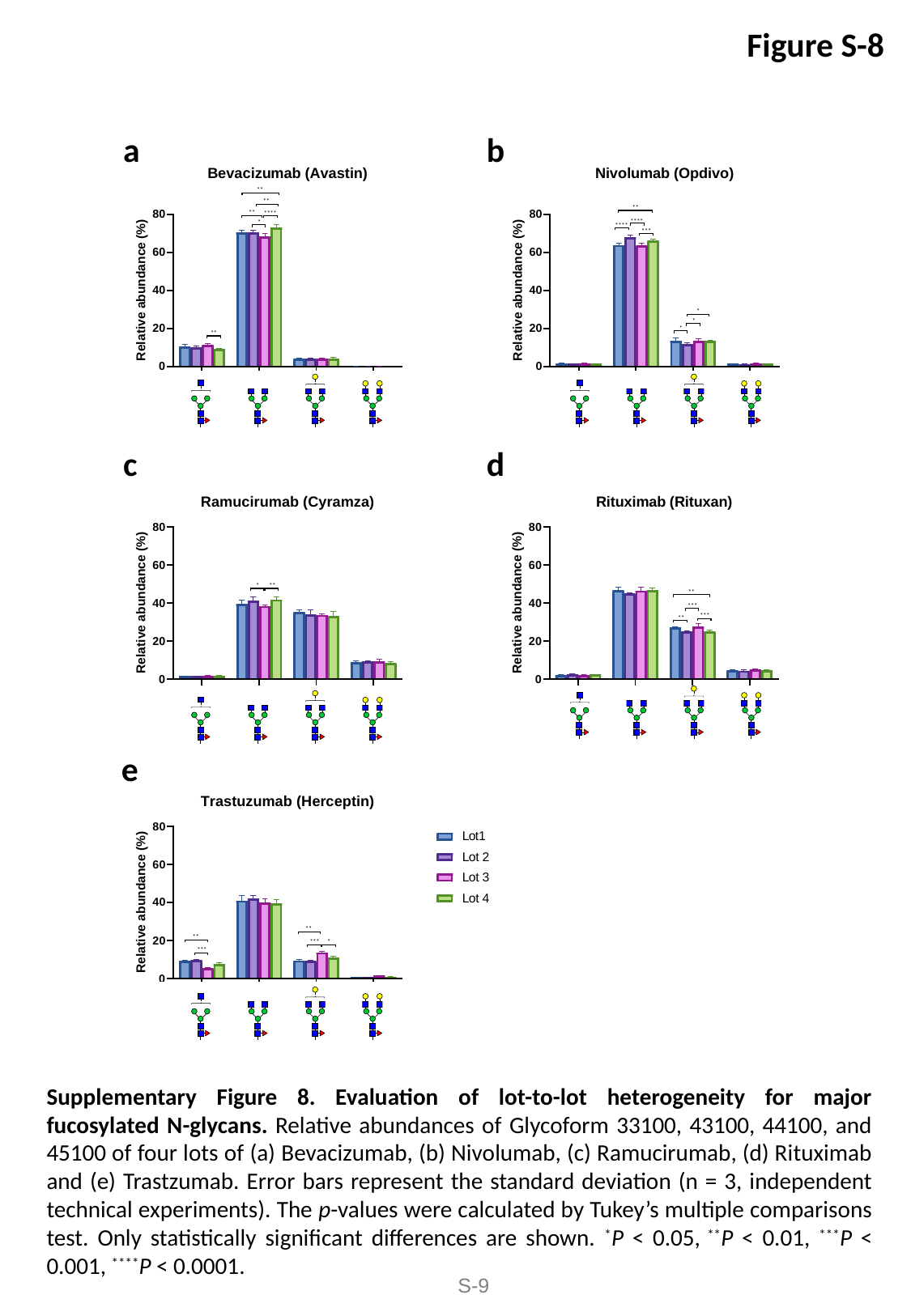

Figure S-8
b
a
**
**
**
**
****
****
*
****
***
*
*
*
**
d
c
**
*
**
***
***
**
e
**
**
*
***
***
Supplementary Figure 8. Evaluation of lot-to-lot heterogeneity for major fucosylated N-glycans. Relative abundances of Glycoform 33100, 43100, 44100, and 45100 of four lots of (a) Bevacizumab, (b) Nivolumab, (c) Ramucirumab, (d) Rituximab and (e) Trastzumab. Error bars represent the standard deviation (n = 3, independent technical experiments). The p-values were calculated by Tukey’s multiple comparisons test. Only statistically significant differences are shown. ^*^P < 0.05, ^**^P < 0.01, ^***^P < 0.001, ^****^P < 0.0001.
S-9

### Slide 10
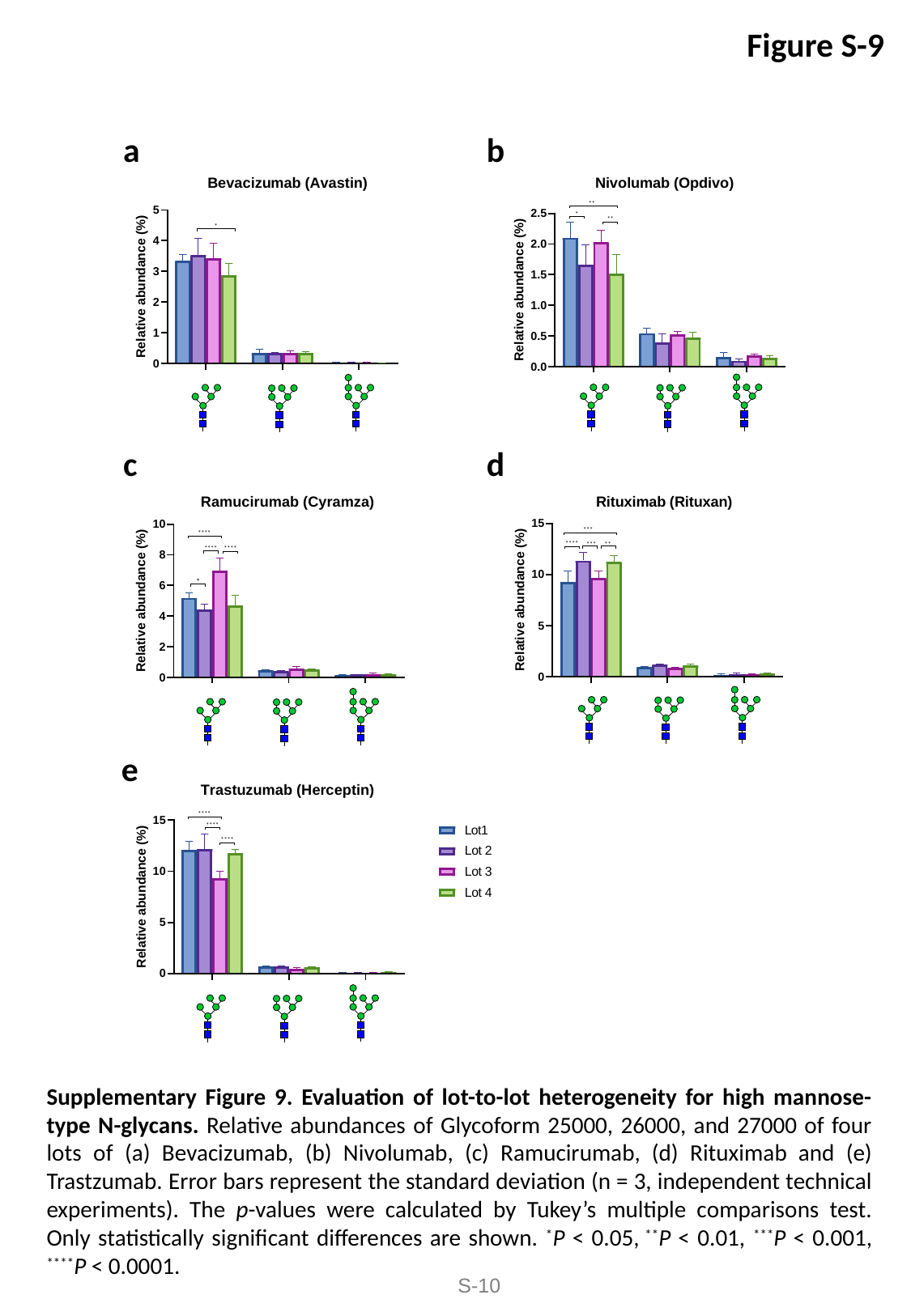

Figure S-9
b
a
**
*
**
*
d
c
***
****
****
***
**
****
****
*
e
****
****
****
Supplementary Figure 9. Evaluation of lot-to-lot heterogeneity for high mannose-type N-glycans. Relative abundances of Glycoform 25000, 26000, and 27000 of four lots of (a) Bevacizumab, (b) Nivolumab, (c) Ramucirumab, (d) Rituximab and (e) Trastzumab. Error bars represent the standard deviation (n = 3, independent technical experiments). The p-values were calculated by Tukey’s multiple comparisons test. Only statistically significant differences are shown. ^*^P < 0.05, ^**^P < 0.01, ^***^P < 0.001, ^****^P < 0.0001.
S-10

### Slide 11
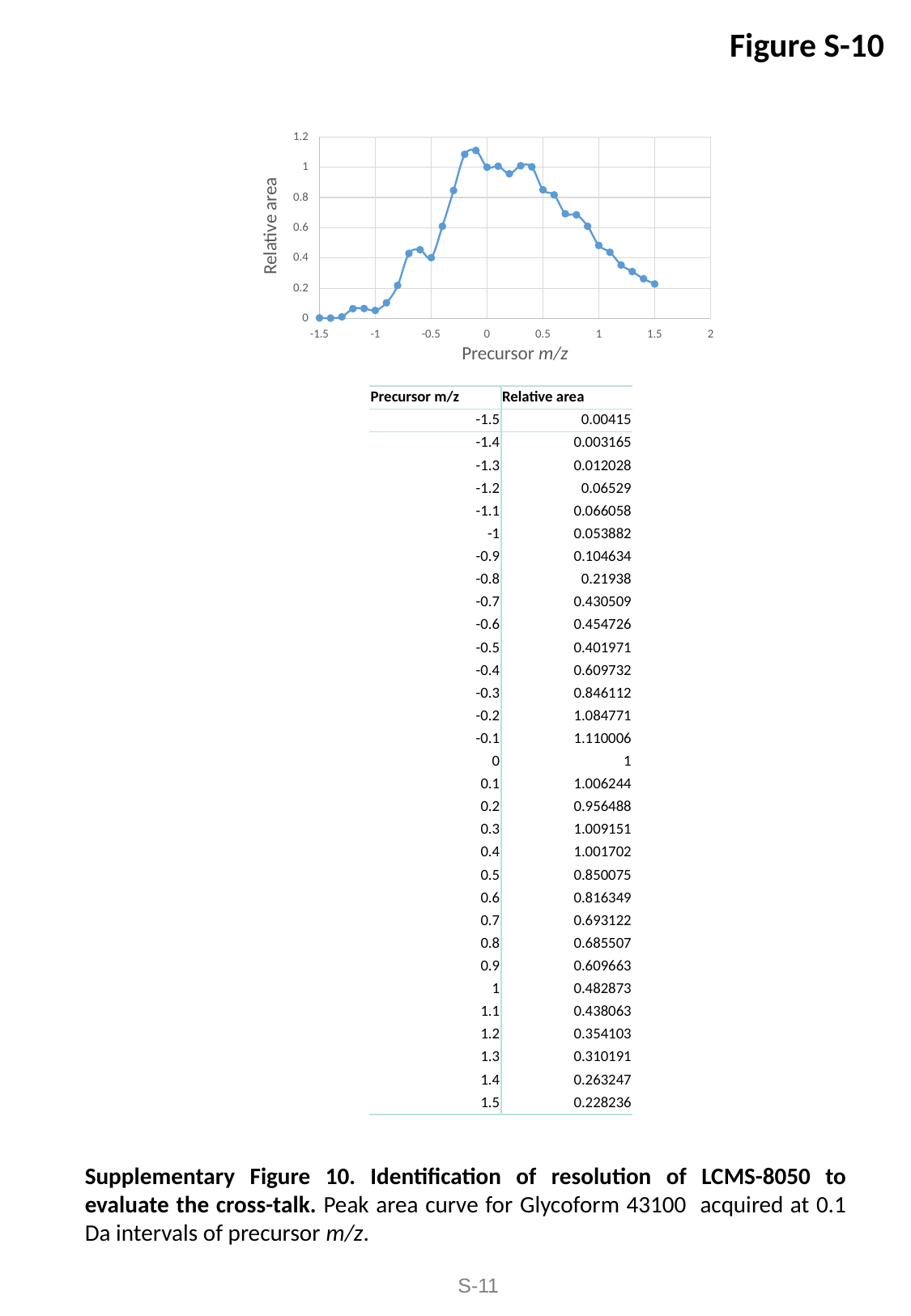

Figure S-10
| Precursor m/z | Relative area |
| --- | --- |
| -1.5 | 0.00415 |
| -1.4 | 0.003165 |
| -1.3 | 0.012028 |
| -1.2 | 0.06529 |
| -1.1 | 0.066058 |
| -1 | 0.053882 |
| -0.9 | 0.104634 |
| -0.8 | 0.21938 |
| -0.7 | 0.430509 |
| -0.6 | 0.454726 |
| -0.5 | 0.401971 |
| -0.4 | 0.609732 |
| -0.3 | 0.846112 |
| -0.2 | 1.084771 |
| -0.1 | 1.110006 |
| 0 | 1 |
| 0.1 | 1.006244 |
| 0.2 | 0.956488 |
| 0.3 | 1.009151 |
| 0.4 | 1.001702 |
| 0.5 | 0.850075 |
| 0.6 | 0.816349 |
| 0.7 | 0.693122 |
| 0.8 | 0.685507 |
| 0.9 | 0.609663 |
| 1 | 0.482873 |
| 1.1 | 0.438063 |
| 1.2 | 0.354103 |
| 1.3 | 0.310191 |
| 1.4 | 0.263247 |
| 1.5 | 0.228236 |
Supplementary Figure 10. Identification of resolution of LCMS-8050 to evaluate the cross-talk. Peak area curve for Glycoform 43100 acquired at 0.1 Da intervals of precursor m/z.
S-11
